# The *de novo* reference genome and transcriptome assemblies of the wild tomato species *Solanum chilense*

**DOI:** 10.1101/612085

**Authors:** Remco Stam, Tetyana Nosenko, Anja C. Hörger, Wolfgang Stephan, Michael Seidel, José M.M. Kuhn, Georg Haberer, Aurelien Tellier

## Abstract

**Background:** Wild tomato species, like *Solanum chilense*, are important germplasm resources for enhanced biotic and abiotic stress resistance in tomato breeding. In addition, *S. chilense* serves as a model system to study adaptation of plants to drought and to investigate the evolution of seed banks. However to date, the absence of a well annotated reference genome in this compulsory outcrossing, very diverse species limits in-depth studies on the genes involved.

**Findings:** We generated ∼134 Gb of DNA and 157 Gb of RNA sequence data of *S chilense*, which yielded a draft genome with an estimated length of 914 Mb in total encoding 25,885 high-confidence (hc) predicted gene models, which show homology to known protein-coding genes of other tomato species. Approximately 71% (18,290) of the hc gene models are additionally supported by RNA-seq data derived from leaf tissue samples. A benchmarking with Universal Single-Copy Orthologs (BUSCO) analysis of predicted gene models retrieved 93.3% BUSCO genes, which is in the current range of high-quality genomes for non-inbred plants. To further verify the genome annotation completeness and accuracy, we manually inspected the NLR resistance gene family and assessed its assembly quality. We revealed the existence of unique gene families of NLRs to *S. chilense*. Comparative genomics analyses of *S. chilense*, cultivated tomato *S. lycopersicum* and its wild relative *S. pennellii* revealed similar levels of highly syntenic gene clusters between the three species.

**Conclusions:** We generated the first genome and transcriptome sequence assembly for the wild tomato species *Solanum chilense* and demonstrated its value in comparative genomics analyses. We make these genomes available for the scientific community as an important resource for studies on adaptation to biotic and abiotic stress in *Solanaceae*, on evolution of self-incompatibility, and for tomato breeding.

## INTRODUCTION

Tomato (*Solanum lycopersicum*) is arguably the most important vegetable crop and an important model organism for fleshy fruit development [1,2]. Together with its wild relatives it is also an interesting model sytstem regarding tolerance to abiotic and biotic stresses such as pathogens. As with many crops, tomato breeders have often used germplasm of wild relatives to improve cultivar quality, including enhanced stress tolerance [3]. Several wild tomato species have been sequenced. Genome assemblies exist, for *S. habrochaites, S. pimpinellifolium* and *S. pennellii.* Yet, fully accessible and annotated reference genomes sequences to date are only available for the cultivated tomato *S. lycopersicum* [1] and the selfing wild tomato relative *S. pennellii* [3] Here we present a reference genome assembly, annotation and additional *de novo* leaf transcriptome assemblies for a stress tolerant and outcrossing wild tomato species, *S. chilense*.

*S. chilense* occurs on the southern edge of the wild tomato species range, in southern Peru and northern Chile. It belongs to the section Peruvianum, which contains four closely related wild tomato species, of which *S. chilense* forms a monophyletic subclade [4]. *S. chilense* split from its nearest sister species *S. peruvianum*, occurring in central and southern Peru, about 1 mya [5,6]. Since then, the species has migrated southward and colonised diverse arid habitats both in mountainous and coastal terrain bordering the Atacama desert and characterized by low temperature or extreme aridity, respectively [7]. (Figure 1) *S. chilense* has been extensively used as an non-model organism for its interesting ecology and thus several studies focused on drought [8] salt [9,10] and cold tolerance [11], as well as for adaptation to extreme environments [12,13]. Furthermore, as an outcrossing species it has been used to understand the breeding system evolution (self-incompatibility) in the tomato clade [14]. The species is characterized by high levels of genetic diversity [5–7] probably due to existence of seed banking [15]. Besides its role as a study system, *S. chilense* has been used as a resource in tomato breeding. For example, genes from *S. chilense* have been successfully used to enhance resistance to the fungal pathogen *Verticilium dahliae* [16] and to the Tomato Mosaic Virus Y (resistance genes *Ty-1* and *Ty-3*) in *S. lycopersicum* [17].

**Figure 1.**
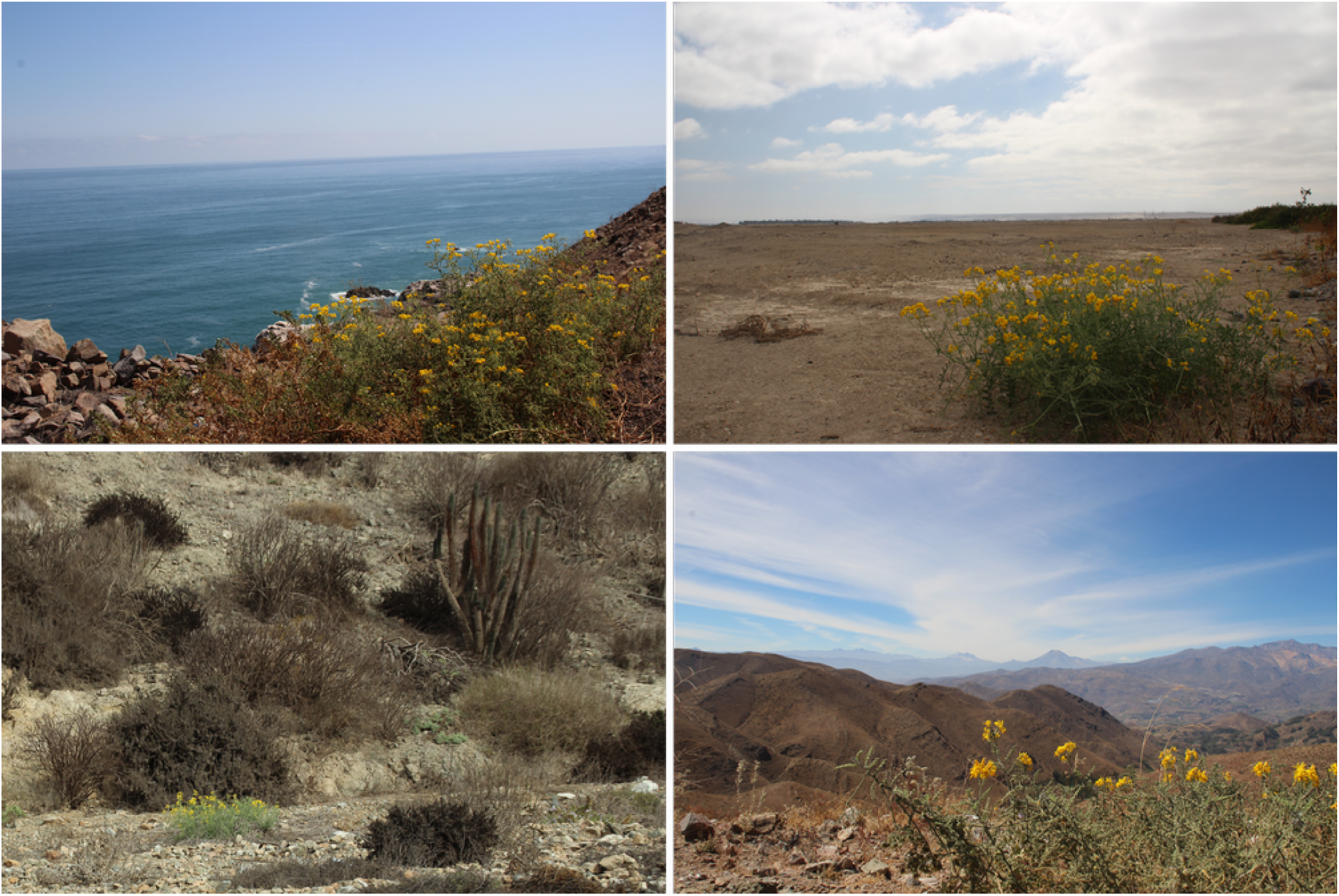
Pictures of *S. chilense* populations in their natural habitat (taken by R. Stam). The top panels show coastal and lowland habitats, the lower panels, typical mountain habitats. LA3111 originates from a mountainous habitat, similar to the last panel.

To corroborate the quality of our reference genome, and to demonstrate its value for future molecular and genomic studies, we compared the NLR family in *S. chilense* with those in cultivated tomato (*S. lycopersicum*) and the wild relative *S. pennellii*. Canonical pathogen resistance genes in plants often belong to the NLR family (Nod-like receptor or Nucleotide binding site, leucine rich repeat containing receptor) [18]. NLRs are modular and contain an N-terminal domain that can be a Toll-Interleukin Receptor (TIR) or a Coiled Coil (CC) domain, followed by a Nucleotide Binding Site (NBS) domain and several Leucine Rich Repeats (LRR). Complete NLRs have all three domains, whereas partial NLRs lack one or the other. NLRs are involved in signalling of the plant immune system and, interestingly, also partial NLRs can be functional in resistance signalling [19]. TIR-domain-containing NLRs are called TNL and CC-domain-containing NLR are referred to as CNL. The latter can again be subdivided into several clades. NLRs are thus divided into several functional sub-clades, for most of which the molecular function is still unknown.

Because of their importance to plant health, NLR evolution has been extensively studied in numerous plant species. Comparative studies in *S. lycopersicum* and some of its wild relatives revealed interesting interspecific differences of the NLR complement [20]. The cultivated tomato and its most closely related relative, *Solanum pimpinellifolium*, contain respectively 326 and 355 NLRs, while *S. pennellii* contains only 216 putative NLRs [21]. These substantial differences in NLR repertoire are hyothesised to be the result of a birth and death process [22] could possibly be explained by differences in pathogen pressure. *S. pimpinellifolium* and ancestors of *S. lycopersicum* are found in northern South-America and Central America in climatic areas possibly more pervasive for pathogens. In contrast, *S. pennellii* for example occurs in generally dryer habitats with lower pathogen pressure, than the cultivated tomato ancestor. Nevertheless, the same functional subclades could be found in these three tomato species, albeit exhibiting different numbers of gene members.

## Data description

### First *S. chilense* genome sequence assembly

Species within the Peruvianum group have diverged relatively recently [4] and exhibit high intraspecific genetic and phenotypic diversity. Hence, species assignment of individuals from this complex can be ambiguous [23]. To confirm that our newly sequenced plant is indeed *S. chilense* we performed phylogenetic comparisons of our sequenced individual and publicly available sequence data from *S. chilense* and *S. peruvianum*. We mapped our sequence data as well as data from all nine publicly available *S. peruvianum* and presumed *S. chilense* data [2,24] (accessions described in Figure 2) against the *S. pennellii* reference genome [3] using STAMPY [25] (substitution rate 0.01, insert size 500). The SNP calling and filtering was done using samtools (mpileup, call -m with default parameters). For all 12 accessions we extracted the sequence at six CT loci (CT066, CT093, CT166, CT179, CT198, CT268). These are single-copy cDNA markers developed and mapped in Tanksley et al. [26] and have previously been used to investigate the evolutionary relationships of wild tomato species (e.g. [6,7,27]). To account for heterozygosity, two alleles were constructed randomly per individual. A concatenated alignment was prepared and manually checked. To this alignment we added 53 sequences obtained by Sanger sequencing in previous work on *S. chilense* and *S. peruvianum* [5]. These sequences originate from *S. chilense* or *S. peruvianum* accessions as identified by the TGRC (UC Davis, USA) according to the taxonomic key in Peralta et al. [28]. *S. ochranthum* (accession LA2682) was used as an outgroup. The phylogentic reconstruction (Figure 2A) was obtained by the Maximum Likelihood method (GTR+Gamm+I algorithm with 1000 bootstrap replicates) as implemented in RaxML [29]. We find that all previously robustly assigned *S. chilense* accessions [5] and our LA3111 individual cluster together into a well-supported monophyletic group (Figure 2A), while the recently sequenced accessions from Aflitos et al [24] and Lin at al [2] form a polyphyleticgroup with known *S. peruvianum* samples. Similar results were were obtained using UPMGA and the Maximum Likelihood method (implemented in Geneious 8) [30].

**Figure 2.**
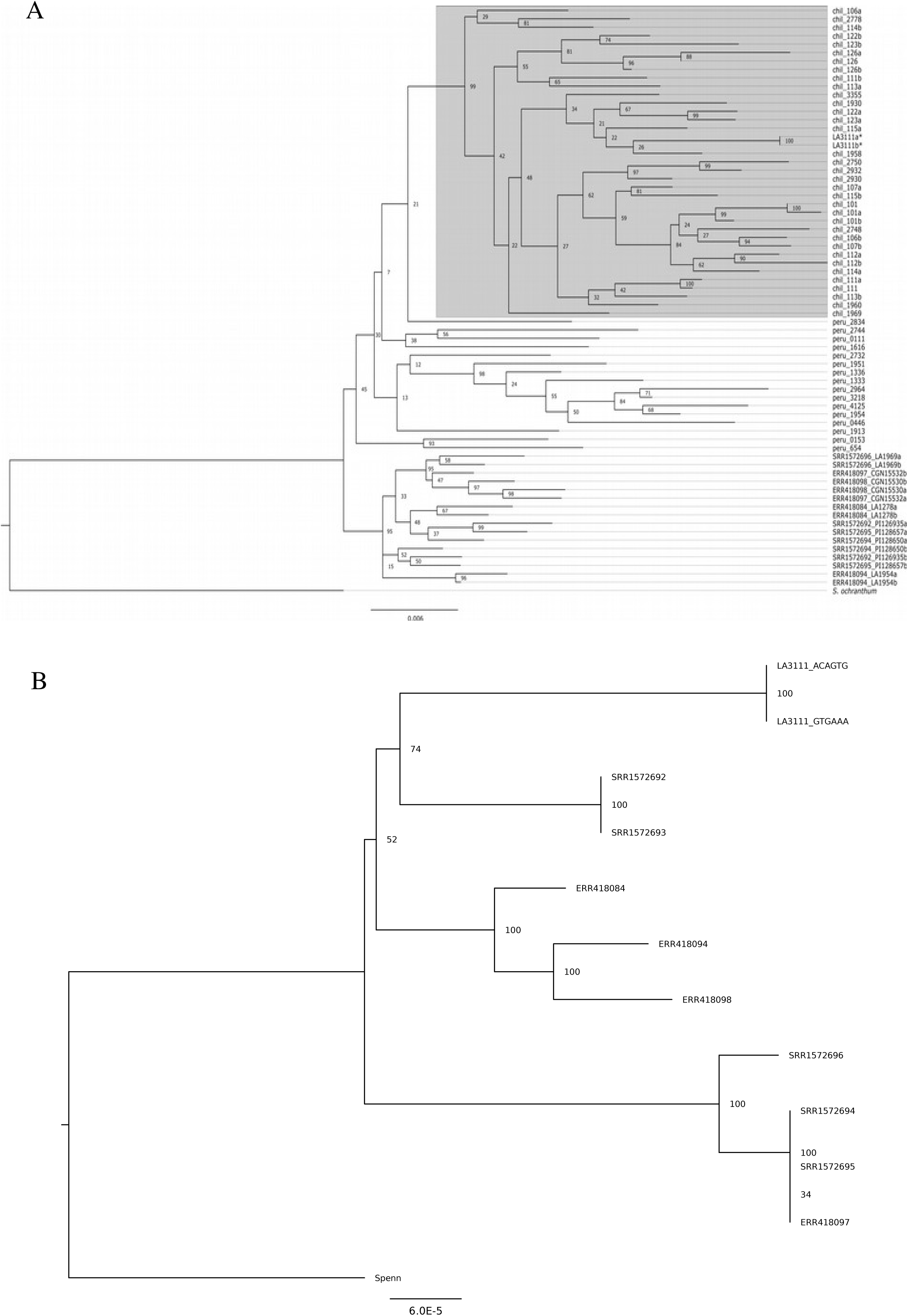
A) Phylogeny based on six CT loci (nuclear genes) extracted from our sequenced *S. chilense* sample and previously sequenced *S. peruvianum* and alleged *S. chilense* samples. Our specimen from accession LA3111, is indicated with *. The phylogeny was constructed after extracting the data mapped to the *S. pennellii* reference genome. A tree was built for the aligned and concatenated sequences using the Maximum Likelihood method (1000 bootstrap replicates). Bootstrap values are reported on each of the branches. *Solanum ochranthum* was used as an outgroup. Chil and peru indicates Sanger sequence from *S. chilense* and *S. peruvianum* individuals, respectively. B) Phylogeny of SNPs in chloroplasts extracted from our sequenced *S. chilense* sample and previously sequenced *S. peruvianum* and alleged *S. chilense* samples. The tree was constructed after extracting the data mapped to the *S. pennellii* reference genome. A tree was built for the aligned sequences using PhyML (GTR, NNI, BioNJ, 1000 bootstrap replicates). Bootstrap values are reported on each of the branches. Individuals ERR418084 and ERR418094: *S. peruvianum* (data from Aflitos et al. 2014), individuals ERR418097 and ERR418098: formerly labelled as *S. chilense*, but probably different species identity (data from Aflitos et al. 2014). This classification has since been withdrawn from the CGN database. The accompanying pictures on the CGN website are not showing *S. chilense* plants. Individuals SRR1572692, SRR1572694 and SRR1572695: *S. peruvianum* (data from Jin et al. 2014), and individual SRR1572696: was reported as *S. chilense* in the main text of the paper (Jin et al., 2014), but the authors confirm it is *S. peruvianum*, as is written in the supplementary data of their paper that contains all origin data. (data from Jin et al. 2014).

Additionally, we reconstructed the chloroplast phylogeny of the members of the *S. peruvianum* clade. We mapped our newly sequenced reads from LA3111, as well as from all nine publicly available *S. peruvianum* and presumed *S. chilense* data (see above) against the *S. pennellii* reference genome [3] using STAMPY [25] (substitution rate 0.01, insert size 500). The SNP calling and filtering was done using samtools (mpileup, call -m with default parameters) and the reconstructed alternative sequences were extracted from *S. pennellii* for the coding regions of the chloroplast for each of the samples. These aligned sequences were used for phylogenetic tree construction using PhyML [31] (ML, GTR, 1000 bootstraps, Best of NNI&SPR, BioNJ). The resulting tree was visualised in and edited for publication using Figtree [32]. All previously sequenced samples are found as a polyphyletic group, which is a topology known for the species *S. peruvianum*, whereas our *S. chilense* sample forms a separated branch (Figure 2B). Thus phylogenetic analyses of both nuclear- and plastid-encoded genes confirm that data presented in this study are the first instance of the *S. chilense* genome sequence assembly.

### *De novo* genome sequence assembly for *S. chilense* LA3111

Four sequencing libraries were produced for one plant from accession number LA3111 with insert sizes of 300bp and 500-550bp for paired-end sequencing, and 8kb and 20kb for long jumping distance protocols. In total we generated ∼134 Gb of raw data (Table S1). We used the Celera assembler (CAv8.3; https://sourceforge.net/projects/wgs-assembler/files/wgs-assembler/wgs-8.3) employing stitched and unassembled MiSeq read data to generate contigs. The fragment correction module and the bogart unitigger of the Celera assembler was applied with a graph and merge error rate of 5%. Minimal overlap length, overlap and merge error rates were set to 50bp and 6% each, respectively. The final contig assembly comprised 150,750 contigs ranging from 1 to 162kb totalling ∼717.7 Mb of assembled genome sequence with a N50 of 9,755 bp. The resulting contigs were linked to scaffolds by SSPACE using all four available libraries of LA3111 [33]. Scaffolds were further processed by five iterations of GapFiller and corrected by Pilon in full-correction mode [34,35]. The 81,307 final scaffolds span a total size of 914 Mb with a N50 of 70.6 kb (Table 1). To check for genome and assembly completeness, we used D-Genies [36] to create a dotplot of our scaffolds against the *S. pennellii* (Figure 3) or *S. lycopersicum* (Figure S1). chromosomes. In both cases, these plots reveal nearly full coverage of the chromosomes compared to *S. pennellii* and *S. lycopersicum*.

**Table 1.** *S. chilense* genome assembly

|  |  |
| --- | --- |
| Total size (Mbp) | 913.89 |
| Scaffolds | 81,307 |
| N50 Scaffolds (bp) | 70,632 |
| Max Scaffold length (bp) | 1,123,112 |
| High confidence gene loci | 25,885 |

**Figure 3.**
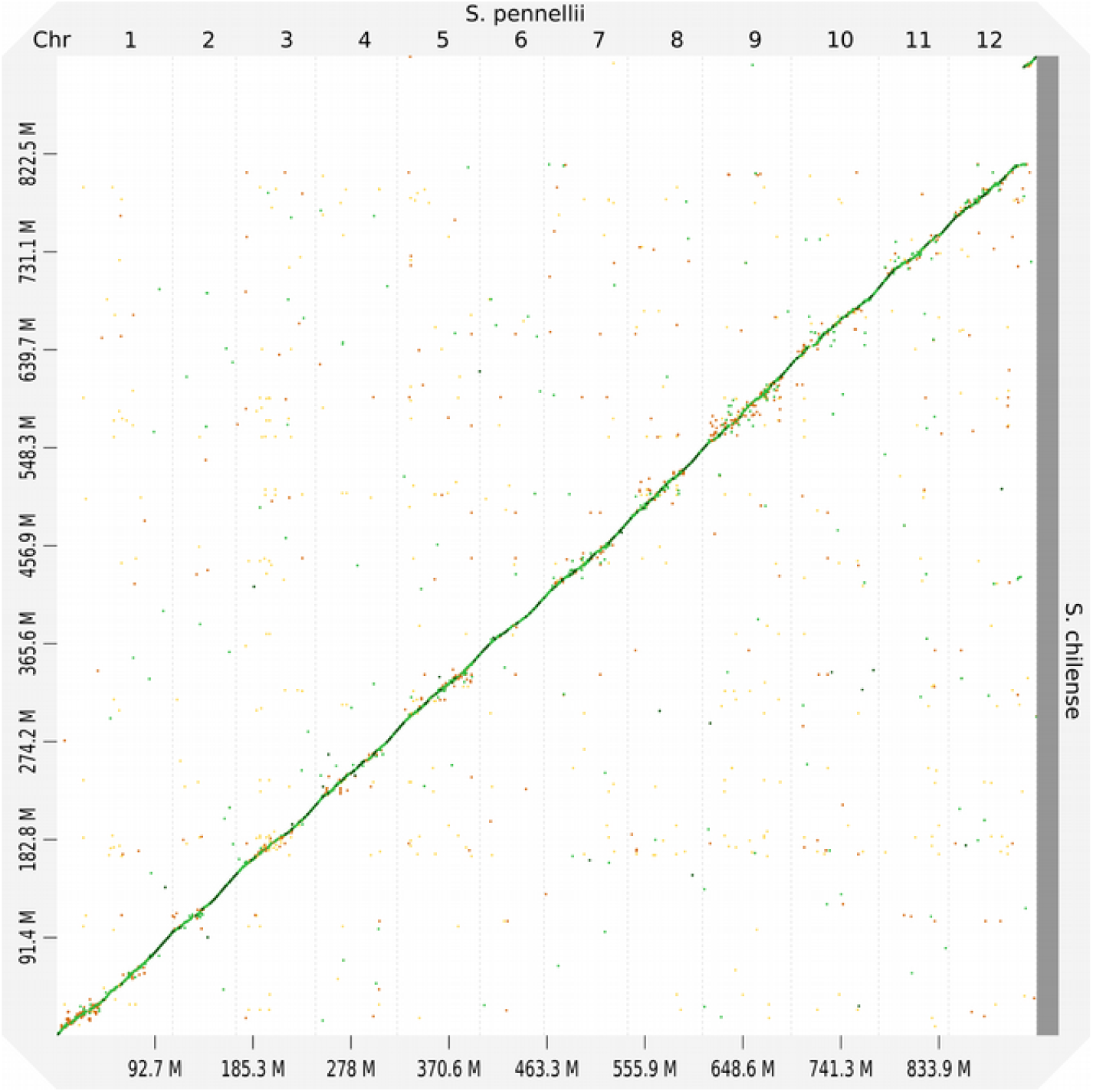
Dotplot analysis of *S. chilense* scaffolds against the *S. pennellii* chromosomes, made using D-Genies [35]. Green lines indicate >75% identity. Orange >60%. The x axis shows the position on *S. pennellii* chromosomes and the y axis on the *S. chilense* scaffolds

### *De novo* assembly of *S. chilense* leaf transcriptome

Twenty four Illumina paired-end read RNA-Seq libraries were generated for 12 *S. chilense* plants from populations LA3111 and LA2750 (Table 2). Replicates were obtained by propagating plants vegetatively. Total RNA was extracted from leaf tissue samples from multiple mature plants under normal and stress (chilling, 6h at 4°C) conditions using the RNeasy Plant Mini Kit (Qiagen GmbH, Hilden, Germany) and purified from DNA using the TURBO DNA-free Kit (Ambion, Darmstadt, Germany). RNA concentration and integrity were assessed using a Bioanalyzer 2100 (Agilent Technologies, Waldbroon, Germany). The preparation of randomly primed paired-end Illumina HiSeq2500 libraries and sequencing were conducted by the GATC Biotech AG. Data for each population were assembled *de novo* using Trinity [37], SOAPdenovo-Trans [38] and Oases-Velvet [39]; the redundancy acquired from pooling the three assemblies was reduced using the EvidentialGene pipeline [40]. The resulting transcriptome assemblies contain 41,666 and 35,470 transcripts and, according to the BUSCO [41] assessement are 93.7 and 94.2% for LA3111 and LA2750, respectively (Table S2).

**Table 2.** *S. chilense de novo* transcriptome assemblies

| Statistics | *S. chilense* transcriptome |  |
| --- | --- | --- |
|  | LA3111 | LA2750 |
| Total contig number | 41,666 | 35,470 |
| Minimum length (bp) | 123 | 123 |
| Maximum length (bp) | 16,476 | 16,473 |
| Average length (bp) | 831 | 943 |
| Median length (bp) | 504 | 684 |
| N50 (bp) | 1383 | 1458 |
| N90 (bp) | 351 | 432 |

### Gene model prediction

We applied a previously described consensus approach [42] to derive gene structures from the *S. chilense* draft genome. Briefly, *de novo* genefinders Augustus [43], Snap [44], and GeneID[45] were trained on a set of high confidence models that were derived from the LA3111 and LA2750 transcriptome assemblies. Existing matrices for eudicots and *S. lycopersicum* were used for predictions with Fgenesh [46] and GlimmerHMM [47], respectively. Predictions were weighted by a decision tree using the JIGSAW software [48]. Spliced alignments of known proteins and *S. chilense* transcripts of this study were generated by the GenomeThreader tool [49]. We used current proteome releases (status of August 2016) of *Arabidopsis thaliana, Medicago truncatula, Ricinus communis, S. lycopersicum, Glycine max, Nicotiana benthiamiana, Cucumis sativa* and *Vitis vinifera*. Spliced alignments required a minimum alignment coverage of 50% and a maximum intron size of 50kb under the *Arabidopsis* splice site model. Next, *de novo* and homology predictions were merged to top-scoring consensus models by their matches to a reference blastp database comprising *Arabidopsis, Medicago* and *S. lycopersicum* proteins. In a last step, we annotated the top-scoring models using the AHRD (“A human readable description”)-pipeline [42] and InterProScan v. 5.21 [50] to identify and remove gene models containing transposon signatures. The resulting final models were then classified into high scoring models according to an alignment consistency of ≥90% for both the *S. chilense* query and a subject protein of a combined *S. lycopersicum* and *S. pennellii* database.

This way, we predicted 25,885 high-confidence (hc) gene loci that show high homology and coverage to known proteins of tomato species. Besides their support by homology, approximately 71% (18,290) of the hc genes are additionally supported by RNA-seq data derived from leaf tissue samples. To obtain the RNA-seq support for the predicted gene models, raw RNA-seq data were processed (adapter and quality trimming) using Trimmomatic v.0.35 [51][Bolger et al., 2014] and aligned to the *S. chilense* genome sequence assembly using STAR v.2.5 [52](Dobin et al. 2013). Read pairs aligned to exonic regions of predicted gene models were summarized per gene using featureCounts [53].

Complementary to the set of hc models, we report the presence of 41,481 low confidence (lc) loci to maximize gene content information. Functionality for some of these models (6,569) is suggested by transcriptome evidence from the leaf RNA-seq data.

Functional gene annotation and assignment to the GO term categories were performed using Blast2GO v. 4.1 [54] based on the results of InterProScan v. 5.21 [50] and BLAST [55] similarity searches against the NCBI non-redundant sequence database. KEGG pathway orthology assignment of protein-coding genes was conducted using KAAS [56].

### Completeness and gene model validation

The completeness of the assembled genome was assessed using BUSCO [41] and was at 91.8% for the genome assembly. Fragments were found for 3.1 additional BUSCO orthologs. These numbers are relatively similar to scores found for previously annotated *S. lycopersicum* and *S. pennellii* (Table S3).

In addition, we assessed synteny between the genomes of three tomato species, *S. chilense* (this study), *S. lycopersicum* (NCBI genome annotation release 102, ITAG2.4), and *S. pennellii* (NCBI genome annotation release 100, v2), Orthologous pairs of protein-coding genes were identified using reciprocal BLAST searches with an e-value threshold of 10^-30^ and maximum target sequence number 50. For *S. lycopersicum* and *S. pennellii*, the longest splice variant for each gene was used as a BLAST input. A spatial distribution of resulting orthologous gene pairs was analysed and gene blocks conserved between genomes (syntenic) were identified using iADHoRe (hybrid mode with minimum syntenic block size = 3; [57]). For tandem arrays of genes, a single representative was retained in syntenic blocks.

We found that our S. *chilense* gene models (hc and lc) show homology to respectively 24,651 *S. lycopersicum* and 25,695 *S. pennellii* genes. Of these, 14,013 and 12,984 genes belong to 2,533 and 2,364 syntenic gene blocks conserved between *S. chilense and S. lycopersicum* or *S. pennellii*, respectively (Table S4, S5). To compare, 977 syntenic gene blocks were detected between *S. lycopersicum* and *S. pennellii* genomes using the same parameters consisting of 18,107 and 17,933 gene models, respectively (Table S6, S7). Synteny dotplots in Figures 3 and S1 illustrate a nearly full coverage between the *S. chilense* scaffolds and *S. pennellii* or *S. lycopersicum* chromosomes. Our gene synteny analyses, confirms that also on gene level our assembly shows large syntenic blocks and thus is relatively complete.

Thus, even though *S. chilense* genome sequence assembly is more fragmented, we can already conclude that the *S. chilense* genome is largely organised as the cultivated tomato and *S. pennellii* genomes, though gene copy numbers vary slightly and small rearrangements did occur.

### NLR identification

To further evaluate the completeness and quality of the *S. chilense* gene model predictions presented in this study, we conducted a detailed analysis of the NLR gene family, a rapidly evolving and thus highly diverse between species group of genes [58]. Loci encoding putative NLR genes were identified using NLRParser [59] with cut-off thresholds as described before [21]. We manually inspected all regions with NLR motifs and updated the annotated open reading frames where this was required. The improved annotation was based on NLR motifs, sequence homology to known NLRs and expression evidence (from the RNA-seq data). In total we found 236 putative NLRs, of which 139 are CNLs and 35 TNLs. 62 NLRs cannot be assigned to either class. Most CDS were supported by all three measures. Only 15 NLR genes were manually curated, using the RNA-Seq data aligned to the reference genome. In ten instances frame shifts made it impossible to enhance the gene model. For these genes the computationally predicted CDS were retained. The remaining 211 predicted NLR gene models showed to be well resolved and did not require any correction. The total number of NLRs identified in *S. chilense the S. chilense genome* is lower than in cultivated tomato (355) and more similar to *Solanum pennellii* (216) (Supplementary material)[3].

The syntenic blocks identified between the *S. chilense* and the *S. lycopersicum* and *S. pennellii* genomes include 69 and 50 hq NLR genes, respectively, and show that NLRs are distributed across all twelve chromosomes (Supplementary material). Except for several short tandems of identical or nearly identical gene copies, NLRs do not tend to form any positional clusters in tomato genomes. Only 30% of *S. chilense* NLRs belong to syntenic gene blocks (compared to *S. lycopersicum* and *S. pennellii*) showing the fast evolution and genomic organisation of this gene family at the phylogenetic time scale (over millions of years).

To further confirm the relative completeness of the NLR set in *S. chilense*, we reconstructed a phylogeny for the gene family based on the NBS protein sequences of the NLRs. Functional clades are assigned based on protein sequences of the NBS, using the same methods as described in Jupe et al. [60]. To define NLR clusters BLASTp searches were used to link new clusters to previously identified ones [60]. In one instance, members of our new cluster matched two previously defined clusters equally well, this cluster thus has double naming (CNL1/CNL9). The NLRs in two identified clusters did not match any NLRs that had been clustered previously, in these cases new cluster numbers were assigned (CNL20, CNL21).

All major NLR clades found in *S. lycopersium* and *S. pennellii* are present in the *S. chilense* genome (Figure 4). There are some small, but interesting differences with other tomato species. The CNL-4 and CNL-15 clusters contained four or five members in *S. lycopersicum*, yet in *S. chilense* each had only one member. In addition, we identified two new clades, CNL20 and CNL21 and when directly comparing *S. pennellii* and *S. chilense*, some clades have more members in the former, and others in the latter (Figure S2) and confirm the birth and death of NLR between species. Similar differences can be seen between *S. pennellii* and *S. lycopersicum* [21].

**Figure 4.**
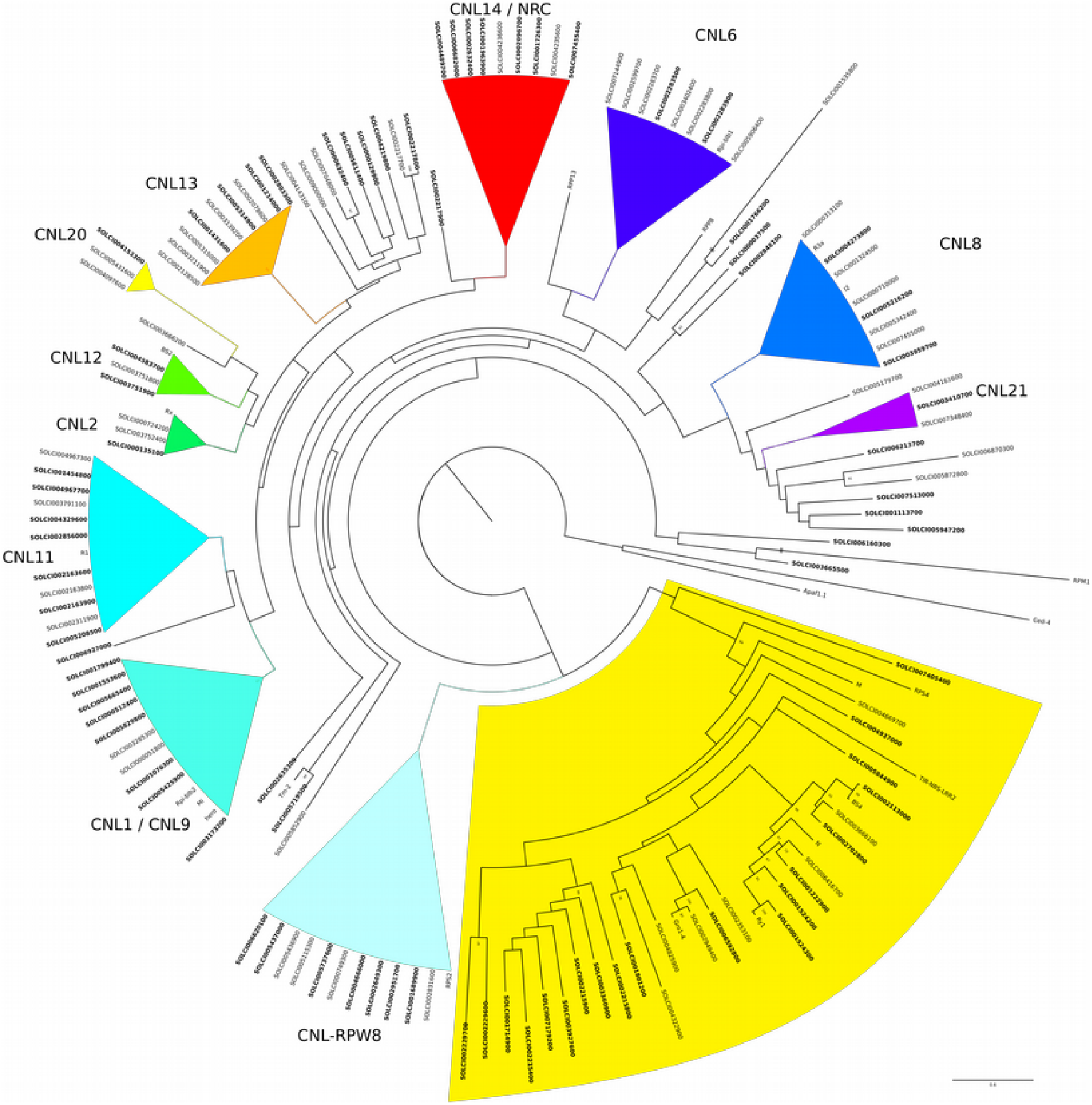
Phylogenetic tree (ML) for the NLRs identified in *S. chilense.* The tree was made as described in Stam et al. 2016 [5]. Clades with high (>80%) bootstrap values are collapsed. Most previously described clades can be identified and are indicated as such. The TNL family is highlighted in yellow. Several previously identified NLR genes from different species are included for comparison and Apaf1.1 and Ced4 are used as an outgroup, similar as in [20,21,59]. NLR with orthologs (based on reciprocal best blast hits) in *S. pennellii* are in bold. Clades CNL20 and CNL21 are new in *S. chilense*.

### Conclusions

We present the draft genome sequence assembly and *de novo* transcriptome assemblies of the wild tomato species *S. chilense.* Using several complementary methods, including comparative analyses for a large and complex gene family such as the NLR-family, we show that quality of this genome assembly and annotation satisfy requirements for a reference genome for comparative genomics studies.

### Data availability

The *S. chilense* genome data and raw RNA-seq data generated for this study deposited to the NCBI Short Read Archive under the BioProject IDs PRJNA508893 and PRJNA474106. The *S. chilense* genome sequence assembly and annotation, CDS and protein models and *de novo* leaf transcriptome assemblies (for the accessions LA3111 and LA2750) are also available as Supplementary Materials and through Sol Genomics Network (https://solgenomics.net/).

## Acknowledgements

RS was supported by the Alexander von Humboldt foundation. *S. chilense* genome sequencing was funded by DFG grant TE 809/7-1 to AT. Generating and sequencing of the *S. chilense* RNA-Seq data was supported by the DFG grant STE 325/15 to WS. We thank the TGRC at UC Davis (USA) for providing the plant material.

## Authors contributions

Conceptualisation: RS, AT, GH, Methodology: RS, GH, TN, AT, Formal analysis: RS, TN, AH, AT, GH,HK, Resources. RS, AH, TN, WS, Data curation: RS, TN, MS, Writing – Original draft: RS, TN, GH Writing – Review & Editing, RS, AH, TN, AT, WS, Visualisation: RS, Supervision RS, GH, AT, WS.

---

**S Figure 1.**
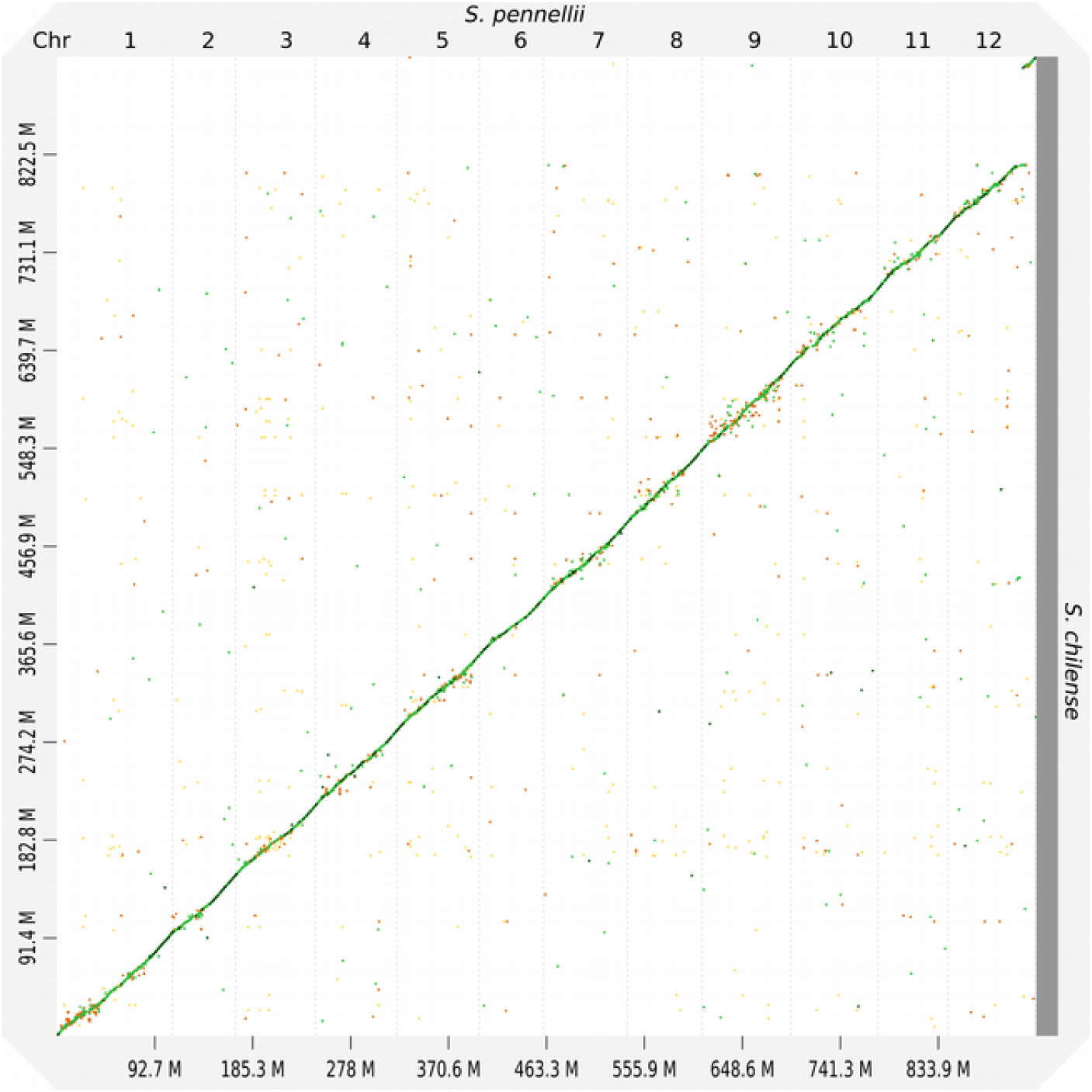
Dotplot analysis of *S. chilense* scaffolds against the *S. lycopersicum* chromosome, made using D-Genies [35]. Green lines indicate >75% identity. Orange >60%. The x axis shows the *S. lycopersicum* chromosomes and the y axis the *S chilense* scaffolds

**S Figure 2.**
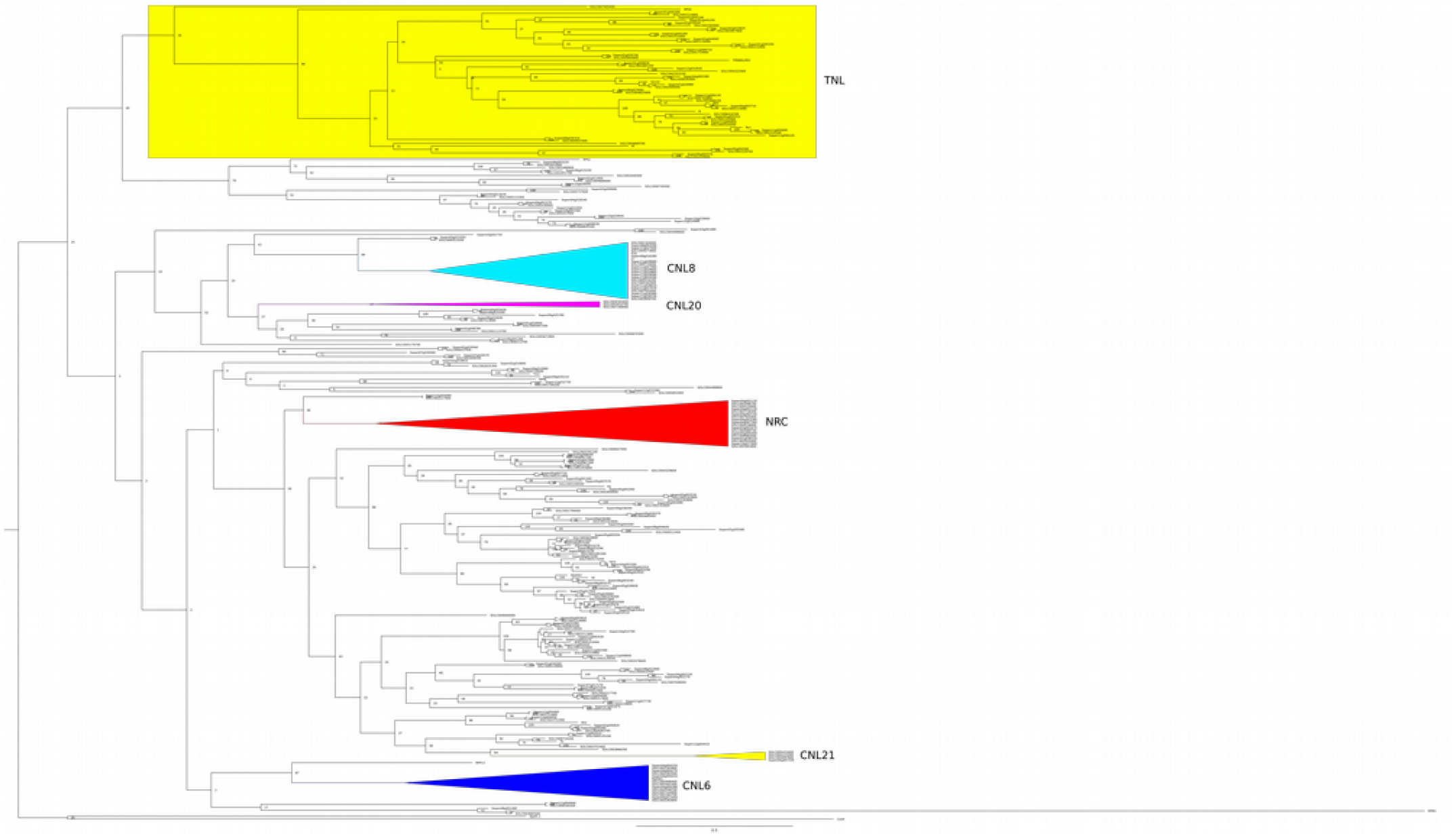
Phylogenetic tree (ML) of *S. pennellii* and *S. chilense* NLRs. Several clades are highlighted to illustrate clades with even numbers (NRC), clades with higher numbers for *S. pennellii* (CNL8), for *S. chilense* (CNL6) and newly discovered clades (CNL20, CNL21)

