## Supplementary figures and images for "The *de novo* reference genome and transcriptome assemblies of the wild tomato species *Solanum chilense*"

### Figure S1.svg.png

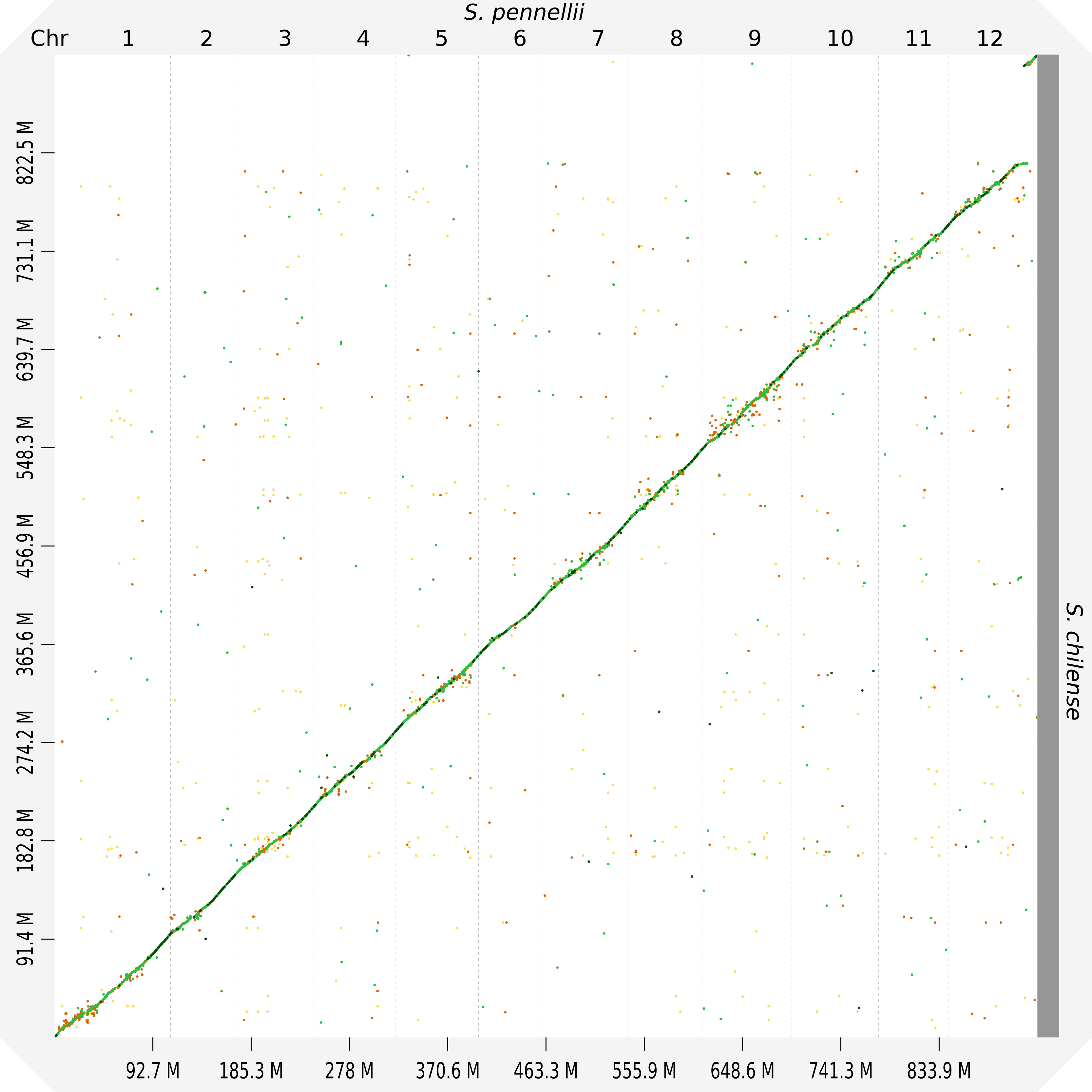

### Figure S2.svg.png

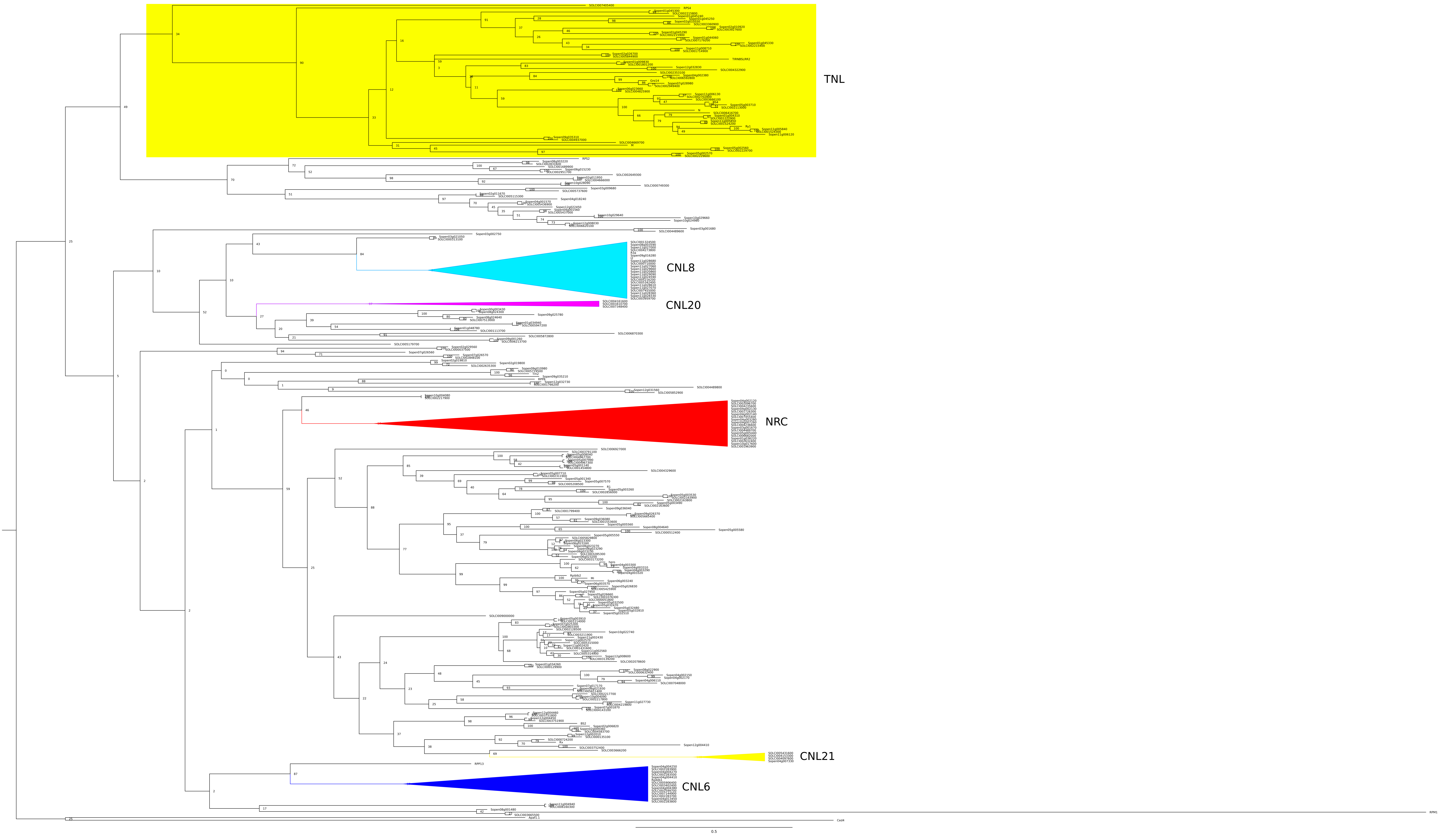
